# Determining the genetic basis of anthracycline-cardiotoxicity by molecular response QTL mapping in induced cardiomyocytes

**DOI:** 10.1101/212381

**Authors:** David A Knowles, Courtney K Burrows, John D Blischak, Kristen M Patterson, Carole Ober, Jonathan K Pritchard, Yoav Gilad

## Abstract

Anthracycline-induced cardiotoxicity (ACT) is a key limiting factor in setting optimal chemotherapy regimes for cancer patients, with almost half of patients expected to ultimately develop congestive heart failure given high drug doses. However, the genetic basis of sensitivity to anthracyclines such as doxorubicin remains unclear. To begin addressing this, we created a panel of iPSC-derived cardiomyocytes from 45 individuals and performed RNA-seq after 24h exposure to varying levels of doxorubicin. The transcriptomic response to doxorubicin is substantial, with the majority of genes being differentially expressed across treatments of different concentrations and over 6000 genes showing evidence of differential splicing. Overall, our observations indicate that splicing fidelity decreases in the presence of doxorubicin. We detect 376 response-expression QTLs and 42 response-splicing QTLs, i.e. genetic variants that modulate the individual transcriptomic response to doxorubicin in terms of expression and splicing changes respectively. We show that inter-individual variation in transcriptional response is predictive of cell damage measured *in vitro* using a cardiac troponin assay, which in turn is shown to be associated with *in vivo* ACT risk. Finally, the molecular QTLs we detected are enriched in lower ACT GWAS *p*-values, further supporting the *in vivo* relevance of our map of genetic regulation of cellular response to anthracyclines.

## Introduction

Anthracyclines, including the prototypical doxorubicin, continue to be used as chemotherapeutic agents treating a wide range of cancers, particularly leukemia, lymphoma, multiple myeloma, breast cancer, and sarcoma. A well-known side-effect of doxorubicin treatment is anthracycline-induced cardiotoxicity (ACT). For some patients ACT manifests as an asymptomatic reduction in cardiac function, as measured by left ventricular ejection fraction (LVEF), but in more extreme cases ACT can lead to congestive heart failure (CHF). The risk of CHF is dosage-dependent: an early study^1^ estimated 3% of patients at 400 mg/m2, 7% of patients at 550 mg/m2, and 18% of patients at 700 mg/m2 develop CHF, where a more recent study puts these numbers at 5%, 26% and 48% respectively^2^. Reduced LVEF shows a similar dosage-dependent pattern, but is not fully predictive of CHF. Perhaps most daunting for patients is that CHF can occur years after treatment: out of 1,807 cancer survivors followed for 7 years in a recent survey a third died of heart diseases compared to 51% of cancer recurrence^3^.

Various candidate gene studies have attempted to find genetic determinants of ACT, but are plagued by small sample sizes and unclear endpoint definitions, resulting in limited replication between studies. Two ACT genome-wide association studies (GWAS) have been published^4^,^5^. While neither found genome-wide significant associations using their discovery cohorts, both found one variant that they were able to replicate in independent cohorts.

A nonsynonymous coding variant, rs2229774, in *RARG* (retinoic acid receptor γ) was found to be associated with pediatric ACT using a Canadian European discovery cohort of 280 patients^4^, and replicated in both a European (*p* = 0.004) and non-European cohort (*p* = 1 × 10^−4^). Modest signal (*p* = 0.076) supporting rs2229774’s association with ACT was also reported in a recent study primarily focused on trastuzumab-related cardiotoxicity^6^. *RARG* negative cell lines have reduced retinoic acid response element (RAREs) activity and reduced suppression of *Top2b*^4^, which has been proposed as a mediator of ACT.

In a different study, a GWAS in 845 patients with European-ancestry from a large adjuvant breast cancer clinical trial, 51 of whom developed CHF, found no variants at genome-wide significance levels^5^. However, one of the most promising variants, rs28714259 (*p* = 9 × 10^−6^ in discovery cohort), was genotyped two further cohorts and showed modest replication (*p* = 0.04, 0.018). rs28714259 falls in a glucocorticoid receptor protein binding peak, which may play a role in cardiac development.

An exciting approach to studying complex phenotypes, including disease, in human is to use induced pluripotent stem cells (iPSC) and derived differentiated cells as *in vitro* model systems for disease. Work by us and others has demonstrated that iPSCs and iPSC-derived cell-types are powerful model systems for understanding cell-type specific genetic regulation of transcription^7^,^8^,^9^,^10^,11, but it is less established whether these systems can be used to model the interplay of genetic and environmental factors in disease progression. Encouragingly, the response of iPSC-derived cardiomyocytes (ICs) to doxorubicin was recently extensively characterized^12^. ICs derived from four individuals who developed ACT after doxorubicin treatment (“DOXTOX” group) and four who did not (“DOX” group), showed clear differences in viability (via apoptosis), metabolism, DNA damage, oxidative stress and mitochondrial function when exposed to doxorubicin. These observations suggest that ICs recapitulate *in vivo* inter-individual differences in doxorubicin sensitivity. Gene expression response differences between the DOX and DOXTOX groups were found using RNA-sequencing data, but the sample size was insufficient (RNA-seq was generated for only 3 individuals in each group) to attempt mapping of genetic variants that might explain the observed functional differences between individuals.

Here we used a panel of iPSC-derived cardiomyocytes from 45 individuals, exposed to five different drug concentrations, to map the genetic basis of inter-individual differences in doxorubicin-sensitivity. We find hundreds of genetics variants that modulate the transcriptomic response, including 42 that act on alternative splicing. We show that the IC transcriptomic response predicts cardiac troponin levels in culture (indicative of cell lysis) in these cell-lines, and that troponin level is itself predictive of ACT. Finally we demonstrated that the mapped genetic variants show significant enrichment in lower ACT GWAS *p*-values.

## Results

### Measuring transcriptomic response to doxorubicin exposure

We generated iPSC-derived cardiomyocytes (ICs) for 45 Hutterite individuals (Figure 1a), and confirmed cardiomyocyte identity (see Methods). We exposed all 45 IC lines to doxorubicin at 5 different concentrations for 24 hours, after which samples were processed for RNA-sequencing. We obtained sufficient read depth (10M exonic reads) for downstream analysis for 217 of the 5 × 45 = 225 individual-concentration pairs, and confirmed sample identity by calling exonic SNPs (see Methods). We observed a strong gene regulatory response to doxorubicin across all concentrations, with 98% (12038 / 12317) of quantifiable genes (5% FDR) showing differential expression across the different treatment concentrations. Our data shows excellent concordance with an existing smaller RNA-seq dataset^12^ (Supplementary Figure S1). Principal component analysis (PCA, Figure 1b) confirms that the main variation in the data is driven by doxorubicin concentration and that the effect of concentration on gene expression is nonlinear. For some individuals the expression data following doxorubicin treatment with 1.25*µM* is closer to the data from treatment with 0.625*µM*, whereas for others it is closer to data from treatment with *2.5µM.* This general pattern provides the first indication in our data that that there is systematic variation in how different individuals respond to doxorubicin exposure. Since the majority of genes appear responsive to doxorubicin we clustered genes into six distinct response patterns using a mixture model approach (Figure 1c, see Methods). From largest to smallest, these clusters represent genes that, through the gradient from low to high concentration treatments, are 1) down regulated 2) initially up-regulated, then further down-regulated 3) up-regulated 4) transiently down regulated 5) transiently up-regulated 6) down-regulated then partially recover (Supplementary Table 1). Gene set enrichments (Supplementary Figure S2, Supplementary Table 2) for the up-regulated cluster include metabolic, mitochrondrial and extracellular processes, as well as known doxorubicin response genes in breast cancer cell lines^13^. The down-regulated cluster shares genes with those down-regulated in response to UV light, which, like doxorubicin, causes DNA-damage. Targets of p53, a transcription factor that responds to DNA damage, are overrepresented in clusters 2 and 5; these clusters involve up-regulation at low concentrations (0.625*µM*) but down-regulation at higher concentrations. Promoter analysis (Supplementary Figure S3, Supplementary Table 3) revealed 21, 45, and 6 significantly enriched transcription factor (TF) binding motifs for clusters 1, 2 and 3 respectively (and none for cluster 4-6). Examples include binding sites for *ZNF145,* a TF that promotes *GPX1* activity and protects cells from oxidative damage during mitochondrial respiratory dysfunction^14^, which is enriched in cluster 1 (down regulation w/ dox); *RONIN,* a regulator of mitochrondrial development and function^15^, which is enriched in clusters 1 and 2; and *MEF2,* myocyte enhancer factor 2, involved in regulating muscle development, stress-response and p38-mediated apoptosis^16^, enriched in cluster 4 (although only at *q* = 0.33).

**Figure 1:**
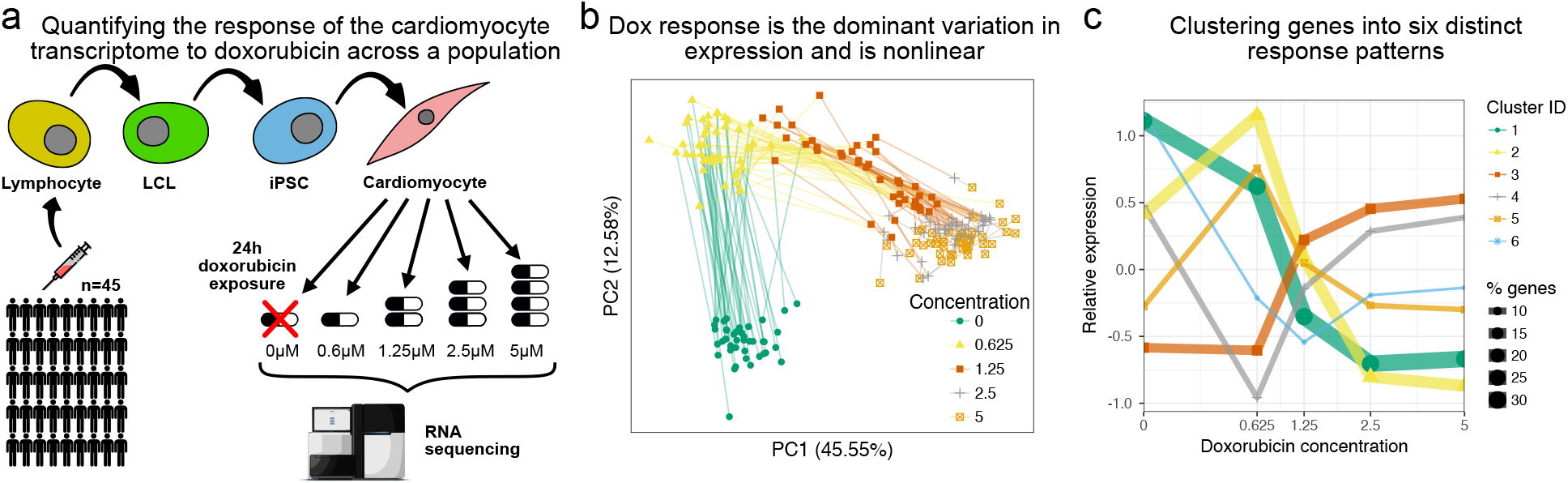
The transcriptomic response of cardiomyocytes to doxorubicin is substantial. a. Cardiomyocytes were derived from lymphoblastoid cell lines (LCLs) of 45 Hutterite individuals, followed by exposure to differing concentrations of doxorubicin and RNA-sequencing. b. PCA of gene expression levels across samples reveals that doxorubicin concentration explains more variance than inter-individual differences, and that the response is non-linear with respect to concentration. c. A probabilistic mixture model uncovers six distinct patterss of response across genes.

### Mapping variants modulating doxorubicin response

We next sought to map single nucleotide polymorphisms (SNPs) that modulate the observed inter-individual transcriptomic response to doxorubicin, leveraging available genetic variation across the 45 individuals^17^. We developed a linear mixed model approach, called `suez` that extends the PANAMA framework^18^ to account for relatedness amongst individuals, repeat measurements, multiple conditions and latent confounding. Testing SNPs within 1Mb of the transcription start site (TSS), 518 genes have a variant with a detectable marginal effect on expression (5% FDR, Supplementary Table 4). Reassuring, these eQTLs show strong replication (Storey’s π_1_ = 0.80) in GTEx heart tissue, and do so more strongly than in GTEx brain (π_1_ = 0.69) or lymphoblastoid cell line (π_1_ = 0.72) data (Figure 2a). Remarkably, even with our moderate number of individuals, we are able to detect many response-eQTLs (reQTLs), i.e. variants that modulate (directly or indirectly) transcriptomic response to doxorubicin. We found reQTLs for 376 genes at a nominal 5% FDR (Supplementary Table 5), which we estimate using a parametric bootstrap corresponds to a true FDR of 8.5% (Figure 2b).

To characterize the detected reQTLs we assigned the response of the major and minor allele to one of the six clusters previously learned (Figure 1c), with heterozygotes expected to display the average of the two homozygous responses. 172 (46%) of reQTLs result in a qualitatively distinct response as determined by the two alleles being assigned to different clusters. The most common transition, occurring for 33 reQTLs, is that the major allele is associated with simple down-regulation (cluster 1) in response to doxorubicin, whereas the minor allele shows a transient up-regulation at low concentration followed by down-regulation at higher concentration (cluster 2).

We further broke-down the significant reQTLs by considering the effect of genotype on expression at each concentration (*β*_*c*_ in Equation 6). We normalized the effect sizes relative to the *β*_*c*_ with the largest absolute value, i.e. we consider 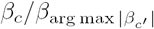, so that the largest genotype effect always corresponds to a normalized value of 1. The resulting normalized effect profiles were split into 9 clusters using *k*-means clustering (Figure 2d). The largest cluster (cluster 1, 85 reQTLs) represents reQTLs with a modest effect size at low concentrations (0,0.625*µM*) which is amplified at higher concentrations (Figure 2e shows a highly significant example). Cluster 2 corresponds to reQTLs whose effect size is attenuated at the 0.625*µM* treatment: examples of reQTLs in this cluster tend to be associated with higher expression level at the 0.625*µM* treatment (e.g. rs16853200’s association with ABCA12 response, Supplementary Figure S4).

**Figure 2:**
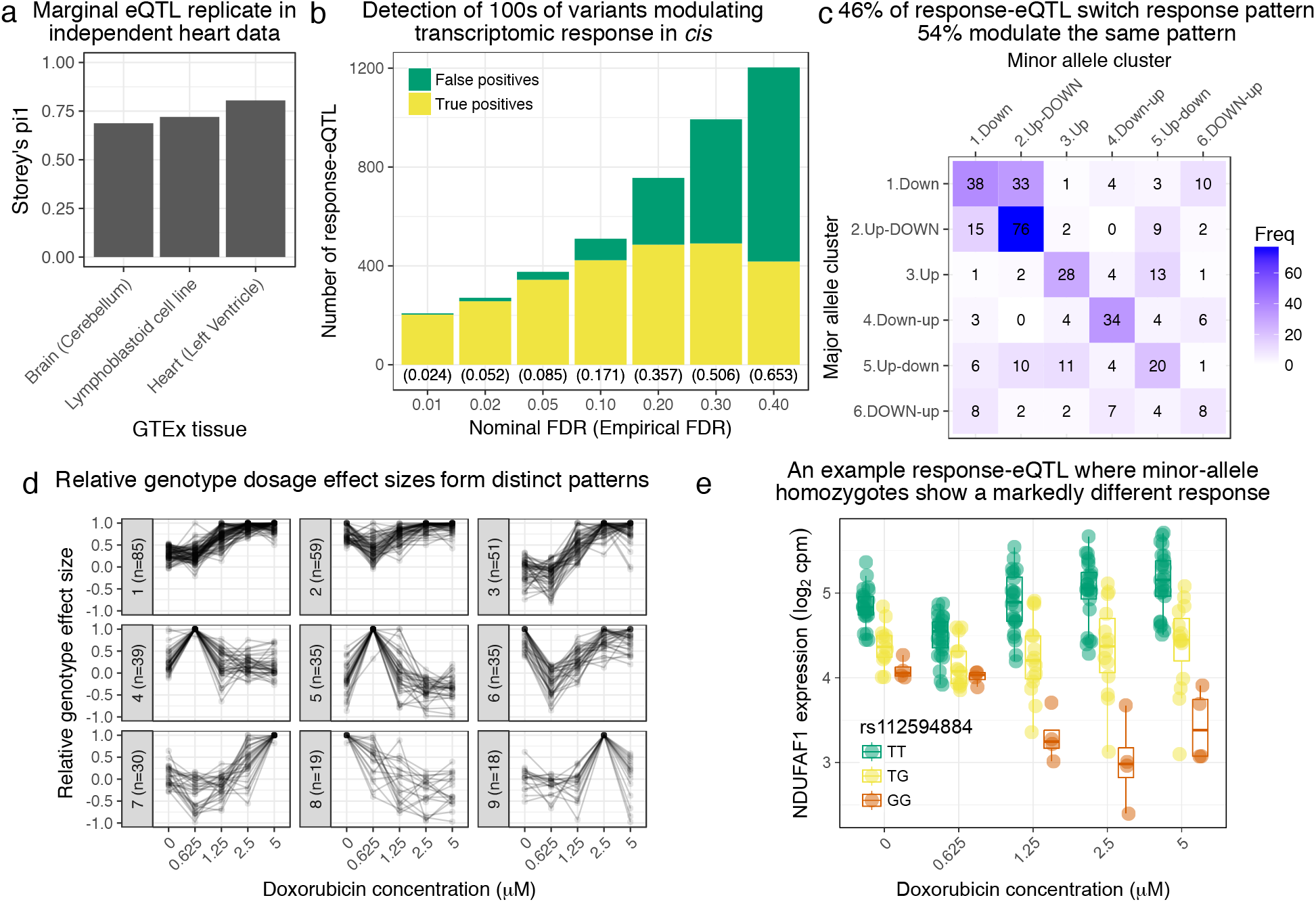
Genetic variation regulates the transcriptomic response to doxorubicin exposure. a. Marginal eQTLs show strong replication in GTEx heart data, and lower replication in other tissues. b. We detect 100s of response-eQTLs (reQTLs): variants that modulate response to doxorubicin. The false positive rate (FPR) is estimated using a parametric bootstrap. c. We developed a statistical method to assign the major and minor allele response to one of the six clusters from Figure 1c. The strongest 46% of detected reQTLs result in a discretely different response, whereas the remainder only modulate the response. d. For significant reQTLs we calculated relative genotype effect sizes by dividing the fitted effect size at each concentration by the (signed) effect size with the largest absolute value. K-means clustering of these effect size profiles reveals distinct patterns, the most common being a small reduction in absolute effect size from 0 to 0.625µM followed by the largest effects being at the highest concentrations. e. An example response-eQTL where rs112594884 regulates the response of the mitochondrial complex I chaperone NDUFAF1. Under the major (T) allele we see moderate down-regulation at 0.625µM followed by up-regulation at higher concentrations. Under the minor (G) allele, there is little change at 0.625µM followed by substantial down-regulation. Since the genotype effects are reduced at 0.625µM and largest at high concentrations this reQTL is assigned to cluster 1 of panel d.

### Doxorubicin exposure reduces splicing fidelity

Oxidative stress, a major downstream consequence of doxorubicin exposure, disrupts splicing of individual genes including *HPRT, POLB*^19^, and *SMA*^20^. We queried the extent to which doxorubicin exposure disrupts splicing patterns across the transcriptome using `LeafCutter`^21^. Across all samples `LeafCutter` detected 27769 alternative splicing “clusters” (referred to here as “ASCs” to avoid confusion with *k*-means clusters), which correspond approximately to splicing events, with a median of 3.0 splice junctions per ASC. Of these, 10430 (59%) ASCs, corresponding to 6398 unique genes, showed an effect of doxorubicin exposure on splicing outcomes (5% FDR, Supplementary Tables 6-7). To characterize these changes we calculated the entropy of the splicing choices made for each significant ASC at each concentration and used *k*-means clusters patterns of change in entropy (Figure 3a). The largest cluster has 6166 ASCs (59%), and corresponds to the null of no clear change in entropy across concentrations. Clusters 2 (*n* = 1136) and 5 (*n* = 475) correspond to increasing entropy with concentration, and clusters 3, 4, 6, 8 and 9 correspond to the maximum entropy being at different concentrations and reaching different maximum levels. Interestingly, only the relatively small cluster 8 (*n* = 304, 3% of ACSs) corresponds to a reduction in entropy at higher concentrations, suggesting the dominant behavior is reduced splicing fidelity and increased alternative splicing in response to doxorubicin.

We further tested the hypothesis that splicing fidelity decreases in the presence of doxorubicin by comparing patterns of intronic percent excised (Ψ) with canonical vs cryptic (unannotated) splice site usage. We clustered the 7792 introns in significantly differentially spliced ASC, that have a change in percent excised (∆Ψ) > 0.1 for some pair of concentrations, into 8 response patterns based on their relative excision proportions across concentrations. For each cluster we calculated the proportion of member introns with neither end annotated, one end unannotated, or both ends annotated (Figure 3b). The clusters representing increased Ψ with concentration (clusters 2, 4, 6 and 7) all show enrichment for cryptic splice site usage. The two most populous clusters (1 and 2) correspond to Ψ decreasing and increasingly continuously with doxorubicin concentration, respectively, and the difference in levels of cryptic splicing is extremely apparent (hypergeometric *p* < 2 × 10^−16^, odds ratio for one annotated end vs two is 28.0).

We additionally used `LeafCutter` quantification of percentage spliced in (PSI) for each splice junction to map splicing QTLs (sQTL) and response-splicing QTLs (rsQTL) using the same methodology as we employed for reQTL mapping. We tested SNPs within 100kb of either end of the splice junction. At 5% FDR we found 467 ASCs with a marginal effect sQTL (Supplementary Table 8) and 42 with a rsQTL (Supplementary Table 9). An example rsQTL is rs72922482’s association with inclusion of exon 2 of APAF1 Interacting Protein (APIP). Under the major T allele exon skipping is extremely rare: the LeafCutter PSI for the spanning junction ranges from 0.00059 to 0.0049 across concentrations (Figure 3c). In rs72922482 heterozygotes, however, the exon is skipped in a significant proportion of transcripts, and this effect is most pronounced in the data collected after treatment at 1.25*µM*, with approximately 50% exon inclusion, suggesting the minor C allele results in very low inclusion of the cassette exon. Another interesting example is *NDUFAF6,* another mitochrondrial Complex I protein, where doxorubicin exposure (particularly at 0.625*µM*) results in increased use of an alternative downstream transcription start site (TSS) which unmasks the influence of rs896853 on a cassette exon between the two alternative TSS (Supplementary Figure S5).

**Figure 3:**
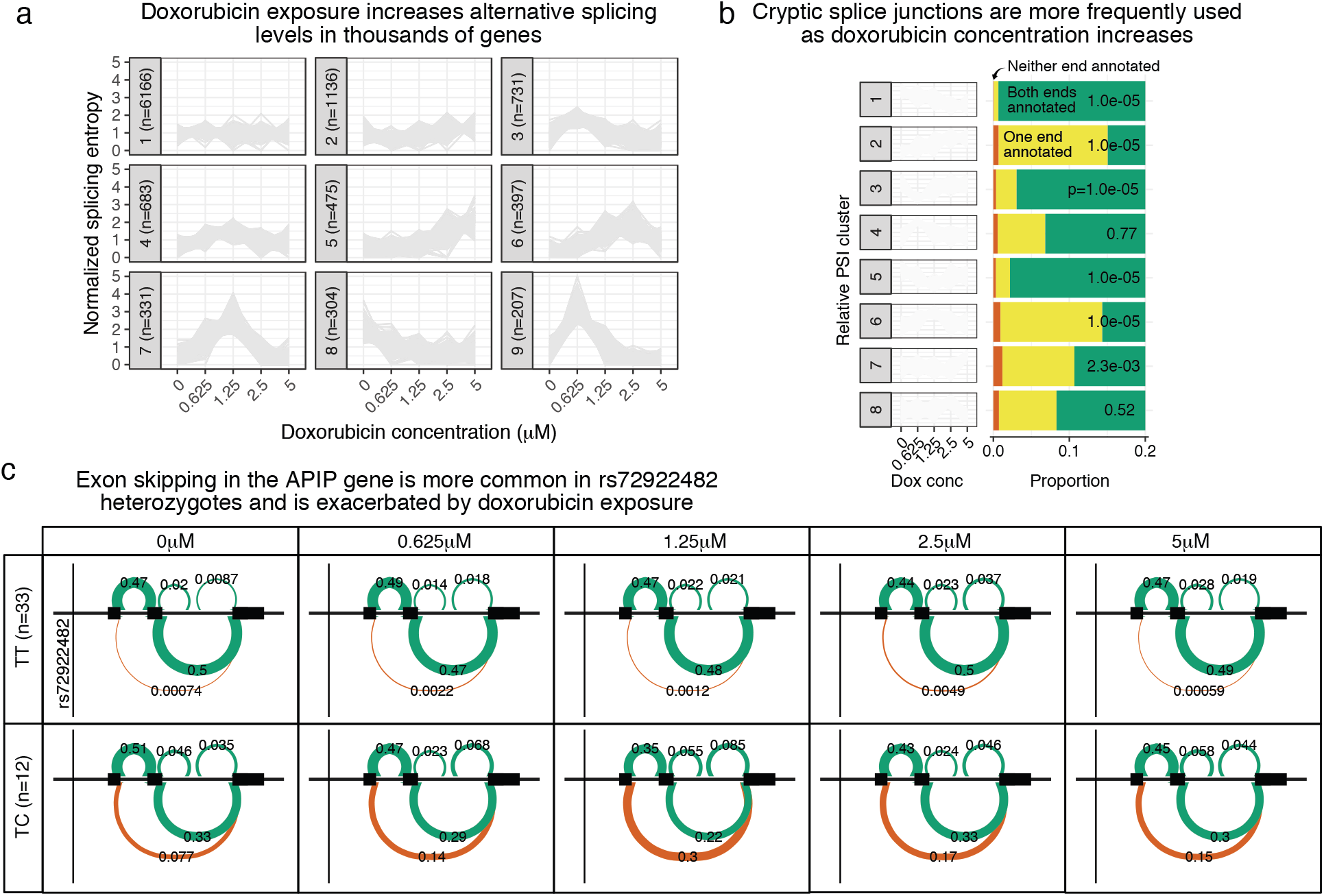
Doxorubicin exposure significantly impacts alternative splicing. a The entropy of splicing choices increases in response to doxorubicin exposure. We measured splicing entropy at different concentrations within `LeafCutter` “Alternative Splicing Clusters” (ACSs) and clustered these into patterns of entropy change. b We separated introns differentially excised with ∆Ψ > 0.1 into 8 clusters based on their relative excision level at each concentration. Introns in clusters corresponding to increased excision at higher doxorubicin concentrations (e.g. cluster 2) are far more likely to use a cryptic (unannotated) splice site at at least one end. p-values shown are for a hypergeometric test of that cluster against all others. c We mapped 42 ASCs with response splicing QTLs, variants that modulate the differential splicing response to doxorubicin.

### Transcriptional response to doxorubicin is predictive of in-vitro cardiac-damage indicator troponin

We used the level of cardiac troponin released into the culture media by lysed cardiomyocytes (see Methods, Supplementary Table 10) to estimate damage occurring as a result of doxorubicin exposure at different concentrations. We observed significant variation in measurable damage caused by doxorubicin across individuals, with 13 of 45 cell lines having a significant correlation between doxorubicin dose and troponin measurement (Figure 4a). We first sought to determine whether the inter-individual variation in troponin in culture could be explained by variation in the overall gene expression response. Since we are interested in this case in inter-individual differences rather than differences between concentration we normalized the troponin measurements to have 0 mean and variance of 1 across samples at each doxorubicin treatment. We found 96.1% (95% credible interval [91.5%98.6%]) or 91.5% of the variance in this normalized troponin level could be explained using gene expression levels (we excluded the troponin genes *TNNT1-3* and *TNNI1-3* from the analysis) at the corresponding doxorubicin concentrations, using a GREML-analysis^22^ or leave-out-one cross validated LASSO^23^ respectively. The optimal LASSO model included 118 genes (Supplementary Table 11).

To further explore the relationship between transcriptomic response and troponin presence in culture, we analyzed differential expression (DE) with respect to troponin measurement at each doxorubicin concentration separately. We found 0, 7, 78, 2984 and 2863 differentially expressed genes (5% FDR, Supplementary Table 12) at the 5 concentrations respectively (Figure 4b). The most strongly DE gene (with respect to effect size) at the 5*µM* treatment is *DUSP* 13, a known regulator of *ASK1* -mediated apoptosis^24^. The large number of DE genes at the 2:5*µM* and 5:0*µM* treatments are broadly shared (nominal replication rate 82% to 85%), and DE genes at the 1:25*µM* treatment generally represent the most strongly DE genes at the higher concentrations (Figure 4c).

**Figure 4:**
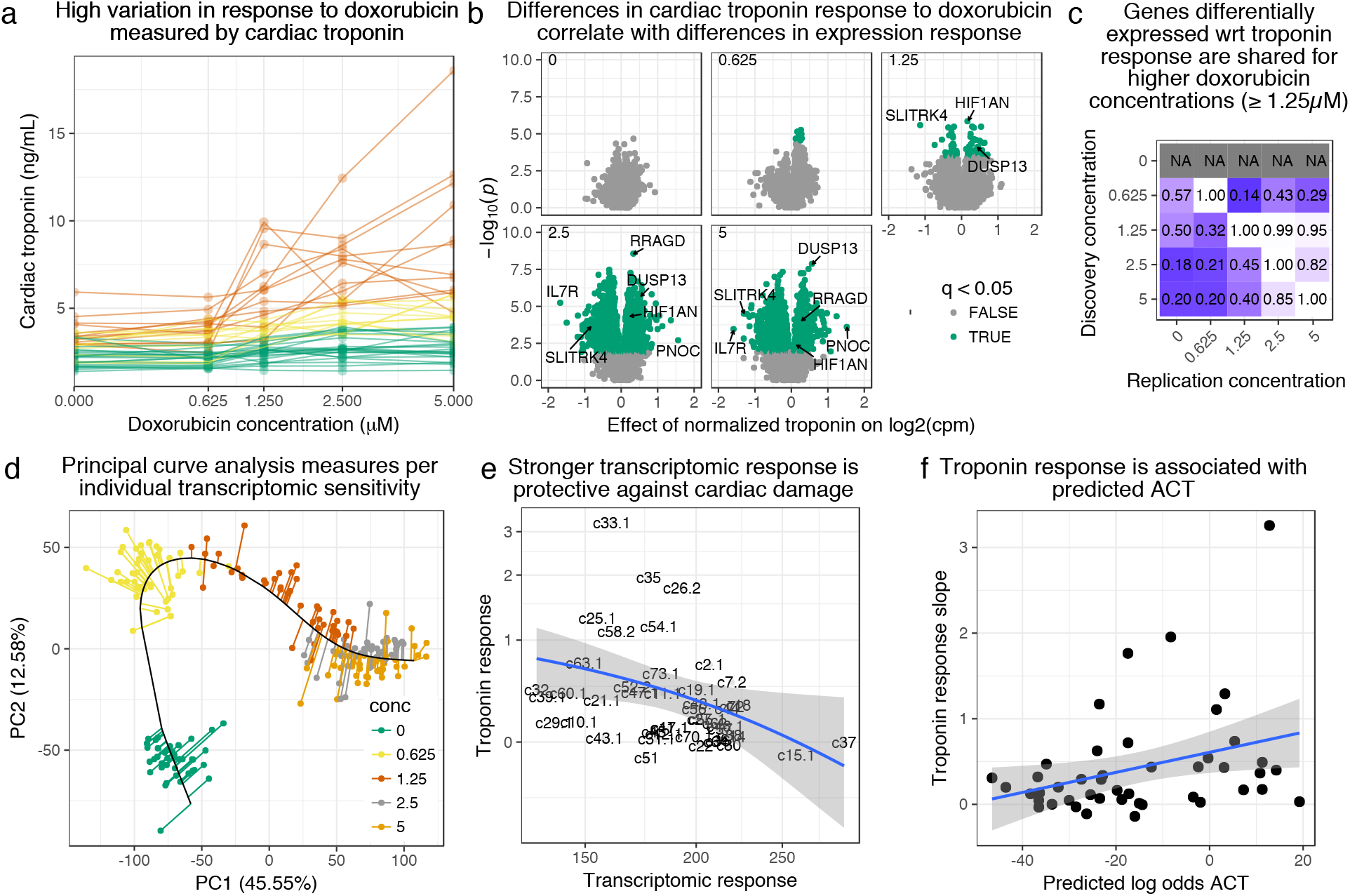
Transcriptomic response is predictive of doxorubicin induced damage as measured by cardiac troponin. **a.** We measured cardiac troponin, a sensitive and specific test for myocardial cell damage, in response to doxorubicin, across all cell lines. **b.** We performed differential expression analyses with respect to troponin at each concentration separately, and observed more differentially expressed genes at higher concentrations corresponding to an increased dynamic range of troponin response. **c.** We took differentially expressed genes (5% FDR) at each concentration and checked for “replication” (nominal p < 0:05) at the other concentrations. Note that no differentially expressed genes were discovered in control condition (0μM). **d.** We summarized gene expression response by first fitting a “principal curve” following increasing doxorubicin concentration, and then measuring the rate of progression along this curve for each individual. **e.** Increased transcriptomic response is associated with reduced cardiac troponin levels, suggesting that the bulk of expression changes we observe are in fact protective against cardiac damage. **f.** We trained a model to predict ACT risk from gene expression response using available 3 v. 3 case/control data^12^ and applied this model to our data. Predicted ACT risk correlated significantly with the slope of troponin response (Spearman ρ = 0:38; p = 0:01), supporting the in vivo disease relevance of our IC system.

To compare troponin measurements to transcriptomic response we determined an overall per-individual level of transcriptomic response with respect to doxorubicin concentration. To this end we fit a principal Curve^25^ through all gene expression samples, initializing the curve to pass sequentially through the successive doxorubicin concentrations (Figure 4d). Projecting every sample on the principal curve gives a single measure of “progression” through response to doxorubicin at increasing concentrations. We then regressed these values against concentration for each individual to obtain a progression rate. We found the troponin measurement slope is significantly negatively correlated (Spearman *ρ* = –0:42; *p* = 0:004, Figure 4e) with the transcriptomic response rate, suggesting that much of the gene expression program being activated in response to doxorubicin is in fact protective against cardiac damage.

Using previously published data^12^, we built a predictive model of ACT risk trained on RNA-seq of ICs exposed to 1*µM* doxorubicin from doxorubicin-treated patients who did (“DOXTOX", *n* = 3) or did not (“DOX”, *n* = 3) develop ACT. Using LASSO with fixed λ = 10^−5^ the optimal model included 17 genes as features (Supplementary Table 13). We applied this model to our expression data from the 0:625*µM* treatment (since this concentration shows excellent concordance with the 1*µM* data of Burridge et al., see Supplementary Figure S1) to obtain predicted log-odds of ACT. While these log-odds are unlikely to be well-calibrated due to differences in the training and test datasets, they may still accurately represent relative risk of ACT across our 45 individuals. Indeed, the log-odds correlated significantly with the troponin response slope (Spearman correlation *p* = 0:01, Figure 4f), suggesting our troponin measurements, and by extension our expression response data, recapitulate in vivo cellular response to doxorubicin.

### Cardiomyocyte molecular QTLs show enrichment in ACT GWAS

To determine the disease-relevance of our molecular QTLs we obtained summary statistics for the largest ACT GWAS to date^5^. While this GWAS was not sufficiently powered to find genome-wide significant associations, 11 variants representing 9 independent loci have *p* < 10^−5^, with the most significant (rs2184559) at *p* = 2:8 × 10^−6^. Of the 8 GWAS variants with *p* < 10^−5^ either tested in our eQTL mapping, or in high LD (*R* ^2^ > 0:8) with a tested SNP, 7 have a nominally significant marginal eQTL (*p* < 0:05, the 8th has *p* = 0:07) and four have a reQTL with *p* < 0:1. The one replicated variant in this GWAS, rs28714259, was not genotyped in our data but is in high LD (*R* ^2^= 0:98) with rs11855704 which is a nominally significant marginal eQTL for tubulin gamma complex associated protein 5 (*TUBGCP5*, Supplementary Figure S6). rs4058287 (GWAS *p*-value 9:68 × 10^−6^) has a marginal effect on Alpha-Protein Kinase 2 (*ALPK2*, also known as “Heart Alpha-Protein Kinase” since it was discovered in mouse heart^26^ and is expressed in few other tissues^27^) expression (*p* = 0:0016) as well as a weak interaction effect (*p* = 0:06, see Figure 5a). Interestingly, *ALPK2* has been shown to upregulate DNA repair genes and to enable caspase-3 cleavage and apoptosis in a colorectal cancer model^28^. The replicating variant from Aminkeng et al. ^4^, rs2229774 only occurs in two individuals in our cohort (who are heterozygous) making eQTL mapping infeasible. Additionally we find a marginal effect eQTL (*p* = 0:0017, Supplementary Figure S7) on SLC28A3 for rs885004, which has previously been associated with ACT in a candidate gene study^29^. rs885004 is intronic, falls in DNase I hypersensitivity and H3K27ac peaks and is in LD (*R* ^2^ = 0:98) with another ACT implicated variant, rs7853758^30^.

**Figure 5:**
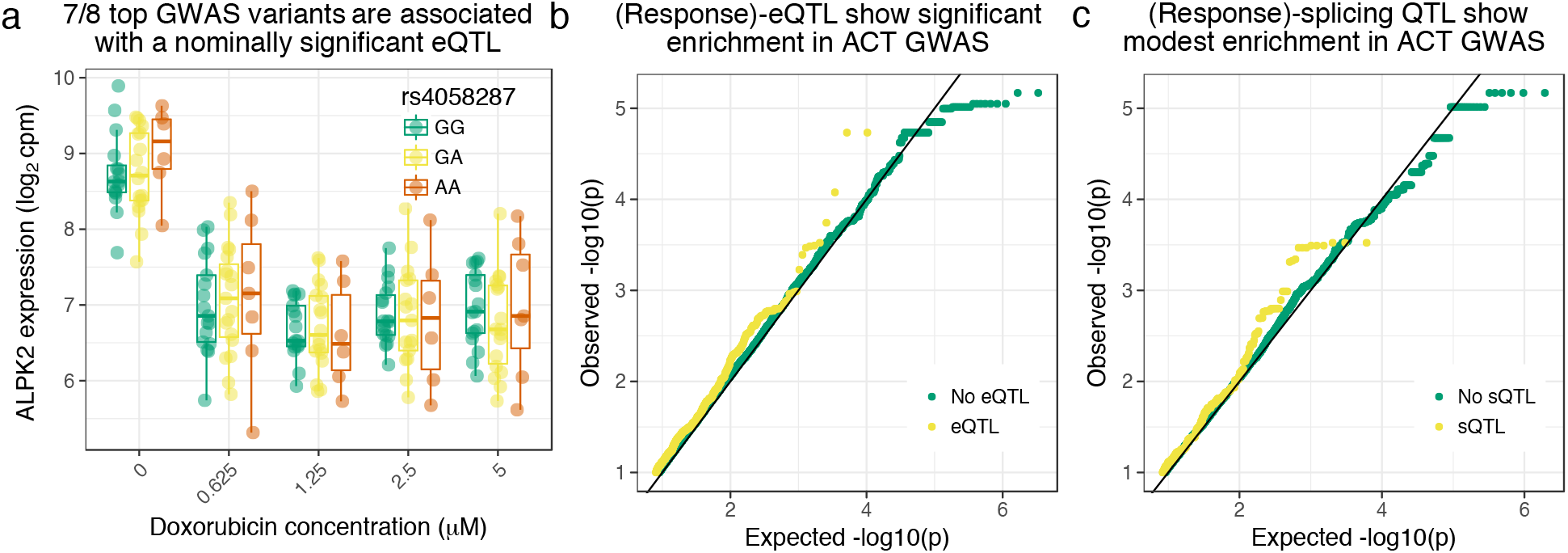
Cardiomyocyte molecular QTLs are enriched in the largest available ACT GWAS^5^. a. rs4058287 has a GWAS p-value of 9:68 × 10^−6^ and is a nominally significant eQTL (*p* = 0:0016) for ALPK2, which is down-regulated in response to doxorubicin. b. SNPs that have either a marginal or response eQTL with *p* < 10^−5^ are enriched in GWAS variants with *p* < 0:05 (hypergeometric test *p* = 3 × 10^−6^). c. SNPs with a marginal or response splicing QTL at *p* < 10^−5^ show modest enrichment in GWAS *p* < 0:005 (hypergeometric *p* = 0:02).

Since the GWAS was overall underpowered, we additionally assessed whether there was detectable enrichment of low GWAS *p*-values for our regulatory QTLs. We considered SNPs with either a marginal or response eQTL with nominal *p* < 10^−5^(corresponding approximately to 5% FDR) and found a significant enrichment for GWAS *p*-values under 0:05 (one-sided hypergeometric *p* = 3 × 10^−6^, OR=1:2, Figure 5b). We observed a more modest enrichment for splicing QTLs: using the same criteria to define a set of sQTL variants we observe significant enrichment in GWAS *p*-values under 0.005 (one-sided hypergeometric *p* =+ 0.02, 5c) but not under 0.05 (one-sided hypergeometric *p* = 0.7).

## Discussion

Human iPSC-derived somatic cells provide a powerful, renewable and reproducible tool for modeling cellular responses to external perturbation *in vitro*, especially for non-blood cell-types such as cardiomyocytes which are extremely challenging to collect and even then are typically only available post-mortem. We established a sufficiently large iPSC panel to effectively query the transcriptomic response of differentiated cardiomyocytes to doxorubicin. We were also able to characterize the role of genetic variation in modulating this response, both in terms of total expression and alternative splicing. There are, of course, caveats associated with using an *in vitro* system, which may not accurately represent certain aspects cardiac response to doxorubicin *in vivo*. That said, the replication of GTEx heart eQTLs, association of troponin levels with predicted ACT-risk^12^, and the observed GWAS enrichment, all support the notion that the IC system recapitulates substantial elements of *in vivo* biology. It is challenging to quantify this agreement however, and there are *in vivo* factors that are certainly not represented: in particular, excessive fibrosis plays a role in ACT^31^, ^32^, ^33^, although it is unclear whether fibroblasts are directly activated by doxorubicin exposure or simply respond indirectly to cardiomyocyte damage.

For many diseases such as ACT which involve an environmental perturbation it is reasonable to suppose that eQTLs detected at steady-state are only tangentially relevant when attempting to interpret disease variants. Such concerns motivated us to focus on response eQTLs, i.e. variants that that have functional consequences under specific cellular conditions because they interact, directly or indirectly, with the treatment. We used a statistical definition of reQTLs corresponding to cases where gene expression levels are significantly better explained using a model including an interaction term between genotype and treatment (represented as a categorical variable), compared to a model with only additive effects for genotype and treatment. Our characterization of the detected reQTL demonstrates that these variants are indeed candidate drivers of differences in individual transcriptomic response to doxorubicin. The strongest reQTL effects correspond to completely different response patterns for the major and minor alleles, while weaker effects correspond to more subtle modulation of the same response pattern. We note that it is not necessarily the case that such reQTLs are the only functionally relevant eQTLs. eSNPs with a marginal (additive) effect on expression of a gene responsive to doxorubicin (as most genes are) could still be important if the relationship between expression and ACT-risk is nonlinear, for example involving thresholding effects.

We observed a statistical enrichment of expression and (to a lesser extent) splicing QTLs in ACT GWAS. However, with no genome-wide significant associations available, fine-mapping of causal variants remains fraught. We anticipate our findings will be increasingly valuable as larger-scale ACT GWAS become available.

We derived ICs from healthy individuals so we do not known which individuals would develop ACT if they required anthracycline treatment. Mapping molecular response QTLs in larger panels of ICs from patients treated with anthracyclines who do or do not develop ACT symptoms would allow stronger conclusions to be drawn about the contribution of the detected (r)eQTLs to disease etiology.

Finally, an interesting observation in our study is that splicing fidelity is reduced upon doxorubicin exposure. This is not completely unexpected since a key downstream side-effect of doxorubicin is increased oxidative stress, which has been previously associated with dysregulated splicing of specific genes^19^,^20^. Our finding that this effect is prevalent across the transcriptome poses further questions about what known effects of doxorubicin might, in fact, be mediated by changes in splicing.

## Methods

### Sample collection and genotyping

Generation of lymphoblastoid cell lines (LCLs) and genome-wide genotyping of many individuals from a multi-generational pedigree were performed previously. Briefly, lymphocytes were isolated from whole blood samples using Ficoll-Paque and immortalized using Epstein Barr Virus^34^,^35^. Phased genotypes were obtained by combining pedigree information, genotypes from SNP arrays, and genotypes from whole genome sequencing of related individuals^17^.

### iPSC reprogramming and cardiomyocyte differentiation

We reprogrammed the 45 LCLs to iPSCs using episomal plasmid vectors, containing *OCT3/4, p53* shRNA, *SOX2, KLF4, L-MYC,* and *LIN28* which avoids integrating additional transgenes^36^. Initially, the lines were generated on mouse embryonic fibroblasts (MEF), which coated the well and served as feeder cells to create an environment supportive of pluripotent stem cells. The colony was then mechanically passaged on MEF and tested for expression of pluripotency-associated markers by immunofluorescence staining and RT-PCR. The lines were passaged for at least 10 weeks on MEF to ensure lines had stabilized. We characterized the iPSC lines using the embryoid body assay, karyotyping, and the PluriTest^37^ classifier. iPSC lines were then transitioned to feeder-free conditions, which was necessary to prime the iPSCs for differentiation. Next we differentiated the iPSCs to cardiomyocytes^38^,^39^. iPSC lines were covered with a 1:60 dilution matrigel overlay for 24 hours. On day 0 iPSC lines were treated with 12*µ*M of the *GSK3* inhibitor, CHIR99021, in RPMI+B27 medium (RPMI1640, 2nM L-glutamine and 1x B27 supplement minus insulin) for 24 hours at which time media was replaced with fresh RPMI+B27. 72 hours after the addition of CHIR99021 (Day 3), 2µM of the Wnt inhibitor Wnt C-59 was added for 48 hours. Fresh RPMI+B27 was added on Days 5, 7 and 10. Beating cells appeared between Days 8-10. These cardiomyocytes consisted of ventricular, atrial and pacemaker-like cells. The cells formed thick layers and contract throughout the well. Metabolic selection was used to purify the cardiomyocytes^40^ from Day 14 to Day 20 when glucose-free RPMI media supplemented with the components essential for cardiomyocyte differentiation^39^, ascorbic acid and human serum albumin, together with lactate, a substrate uniquely metabolized by cardiomyocytes, was added to cells. Because this lactate media can only be metabolized by cardiomyocytes, the non-cardiomyocytes in the culture were removed over the 6 day treatment. On day 20 the cardiomyocytes, now at a high cTnT purity, were replated for experiments in media that contains only galactose and fatty acids as an energy source. This galactose media forces the cardiomyocytes to undergo aerobic respiration, rather than anaerobic glycolysis common in cultured cells.

### Doxorubicin exposure

We incubated the cardiomyocytes in 0, 0.625, 1.25, 2.5, or 5 *µ*M doxorubicin. After 24 hours, we collected the serum and cells from each condition. From the serum, we measured cardiac Troponin T levels using the ABNOVA Troponin I (Human) ELISA kit (cat. no. KA0233). From the cells, we extracted RNA for sequencing. Each treatment batch contained 1 to 4 individuals. RNA quality was assessed with the Agilent Bioanalyzer.

### RNA-sequencing

We prepared libraries using the Illumina TruSeq Library Kit and generated 50bp single-end reads on a HiSeq 4000 at the University of Chicago Functional Genomics Facility. We confirmed sequencing quality using FastQC and MultiQC^41^. We confirmed sample identity by 1) comparing allelic counts (quantified using samtools mpileup^42^) of exonic SNPs to the known genotypes and 2) running verifyBamID^43^.

### Expression quantification

We aligned RNA-seq reads using STAR version 2.5.2a^44^ to GRCh38/GENCODE release 24. We counted reads using featureCounts^45^ and calculated counts per million reads (cpm) using ‘cpm’ from the ‘edgeR’ R package (version 3.18.1)^46^. We discarded samples with < 10^9^ reads and genes with median log_2_(*cpm*) less than 0.

### Differential expression analysis

We performed differential expression (DE) analysis across all 5 doxorubicin concentrations jointly, using either a linear model on quantile normalized cpm value or Spearman correlation, followed by Benjamini-Hochberg False Discovery Rate (FDR) control. Since the vast majority of genes showed differential expression we did not investigate better powered DE methods such as DESeq2.

We clustered genes into “response patterns” using a K-component mixture model

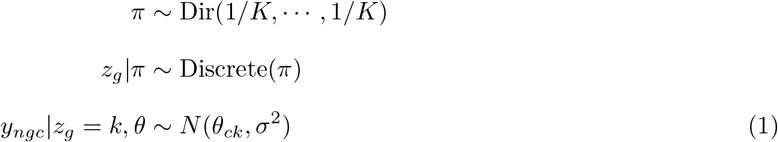

where π is a prior probability vector over cluster assignments, Dir is the Dirichlet distribution, *z*_*g*_ is cluster from which gene *g* is generated, *y*_*ngc*_ is the expression of gene *g* in individual *n* at concentration c, *θ*_*ck*_ is the mixture parameter (mean) across concentrations for cluster *k*, and *σ*^2^ is a shared noise variance. We marginalize (sum) over *z*_*g*_ and optimize with respect to *π, θ, σ* using the rstan R package^47^ (version 2.16.2). The hyperparameters of the Dirichlet distribution are set such that in the limit of large *K* the model approximates a Dirichlet process mixture^48^ which automatically learns of an appropriate number of mixture components to use from data.

Gene set and promoter motif enrichment were performed using HOMER v4.9.1^49^ using default parameters and without *de novo* motif search.

### Response eQTL mapping

We developed an extension of the PANAMA^18^ linear mixed model (LMM) framework to map eQTLs and response eQTLs while accounting for latent confounding, which we call `suez. suez` entails a two step procedure. Step one is used to learn latent factors from all genes, using the model

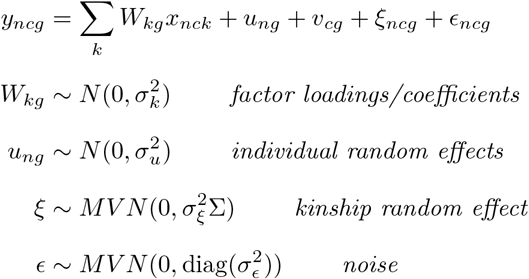

where *x*_*nck*_ are latent factors, *v*_*cg*_ are per gene, per concentration fixed effects. We integrate over *W, u, ξ* and *ε*, which results in a per gene multivariate normal,

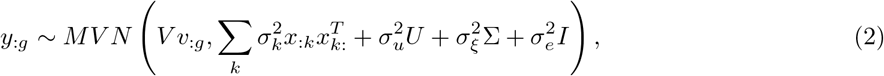

where *y*_*:g*_ refers to the vector of expression for gene *g* across all individuals and concentrations (i.e. all “samples” where a sample is an individual-concentration pair), *V* is a matrix mapping concentrations to samples (i.e. *V*_*sc*_ = 1 iff sample *s* is at concentration *c*) and *U* is a matrix of which samples are for the same individual (i.e. *U*_*ss′*_ = 1 if sample *s* and sample *s′* come from the same individual). We optimize *x,v* and the variances 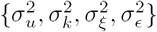 jointly across all genes *g*.

In step 2 we test individual gene-SNP pairs while accounting for confounding using the covariance matrix

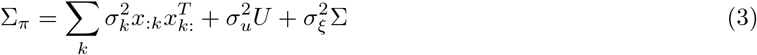

which includes both latent confounding, individual random effects and similarity due to kinship. We consider three LMMs, all with the same parameterization of the covariance 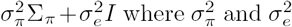 are optimized along with the fixed effects to allow the extent to which each gene follows the global covariance pattern to be adapted. The simple structure of this covariance also allows pre-computation of the eigen-decomposition of Σ_π_ which enables linear (rather than cubic) time evaluation of the likelihood and its gradient.

Model 0 involves no effect of the SNP (and can therefore be fit once for a gene), a fixed effect for concentration. Model 1 adds a marginal effect of the SNP genotype dosage *d*. Finally model 2 adds an interaction effect between concentration and genotype, which is equivalent to a concentration-specific genotype effect.

In summary:

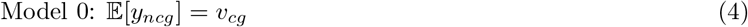

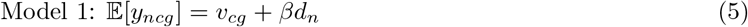

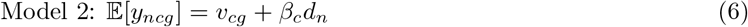

We optimize 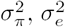 and the regression coefficients for each of the three models separately, and use likelihood ratio tests (LRT) to compare the models. Comparing Model 1 vs 0 (one degree of freedom) tests whether there is a marginal effect of the variant. Comparing Model 2 vs 1 (*C* − 1 = 4 degrees of freedom, where *C* is the number of conditions/concentrations) tests whether there is an interaction effect, i.e. whether the genetic effect on expression is different at different concentrations (or equivalently whether the response to doxorubicin is different for different genotypes). Finally Model 2 vs 0 (*C* = 5 degrees of freedom) tests whether there is any effect of genotype on expression, either in terms of a marginal or concentration-specific effect. We use the conservative approach of using Bonferroni correction across SNPs for a gene, followed by Benjamini-Hochberg FDR control.

We quantile normalize the expression levels across all samples for each gene to a standard normal distribution so that the distributional assumptions of our linear mixed model are reasonable. However, optimizing the variance parameters 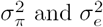 means that the *χ*^2^ distribution for the LRT will only hold asymptotically and p-values for finite sample sizes will tend to be somewhat anti-conservative. To account for this for response-eQTLs, we use a parametric bootstrap since there is no fully valid permutation strategy for testing interaction effects. This involves first fitting Model 1 and then simulating new expression data under the fitted model. Models 1 and 2 are then (re)fit to this data and compared using an LRT. We then perform Bonferroni correction across SNPs for each gene to obtain an empirical null distribution of per gene *p-*values which we use to estimate the true FDR for our response-eQTL results.

For significant reQTLs we assigned the response of the minor allele and major allele to the previously determined clusters using the model

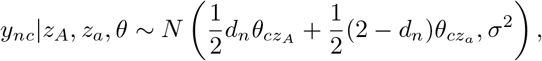

where *y*_*nc*_ is the expression for individual *n* at concentration *c, z*_*A*_ and *z*_*a*_ are the cluster assignments for the major and minor allele respectively, *d*_*n*_ ∈ {0, 1, 2} is the genotype dosage, and *θ* and *σ*^2^ are fixed at the values learned in Equation 1. For each reQTL separately we calculate the likelihood of *y* given all possible pairs of assignments (*z*_*A*_, *z*_*a*_) and choose the maximum likelihood solution.

As for all *k*-means clustering in the paper, we used `KMeans_rcpp` function of the R package `ClusterR` v1.0.6, taking the best of 10 initializations using the `k-means++` option, to cluster the normalized genotype effect profiles of the significant associations. The choice of 9 clusters was determined manually.

### Splicing analysis

We ran `LeafCutter` v0.2.6_dev (using default settings) which allows joint differential intron excision testing across more than two conditions. For each Alternative Splicing Cluster (ASC) `LeafCutter` fits a set of *PercentSplicedIn* probability vectors *ψ*_*c*_, across detected splice junctions *i*, at each concentration c. For ASCs determined to be significantly (5% FDR) differential spliced across concentrations, we calculated the entropy 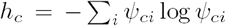 at each concentration *c*. We normalized these profiles as 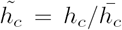 and clustered these profiles, using `KMeans_rcpp` as above.

To investigate the relative usage of cryptic splice sites we first determined the set of 7792 splice junctions that a) fell in ASCs determined to be significantly differentially spliced (5% FDR) and b) had max_*c*_*ψ*_*ci*_ — min_*c*_*ψ*_*ci*_ > 0.1. We obtained normalized intron excision rates by subtracting the per intron mean and dividing by the per intron standard deviation. These *ψ* profiles were clustered using `KMeans_rcpp`. Cryptic splice site usage was determined by considering all exons in Gencode v26 and ignoring transcript structure (i.e. a junction spanning two splice sites used but only in different transcripts would still be considered “annotated”).

For (response) splicing QTL we calculated within ASC intron excision *ψ* with pseudocount of 0.5, and set entries with 0 denominator (no reads for that ASC in that sample) to the mean across all other samples. These values were then 1) *z*-score normalized across samples and 2) quantile normalized to a normal across introns. QTL mapping was then performed using suez considering each intron as a “gene”.

### Modeling cardiac troponin level

We assessed the proportion of variance in cardiac troponin explained by gene expression response. Let *y*_*ci*_ represent the troponin level measured in individual *i* at doxorubicin concentration *c*, normalized to have 0 mean and variance 1 across individuals at each concentration. Let *x*_*cig*_ be the expression of gene *g* (in individual *i* at concentration *c*), *z*-score normalized across samples. We consider the linear model

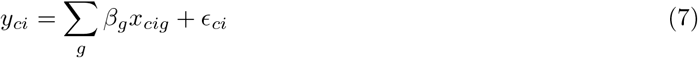

where 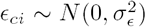 is noise and the coefficients *β*_*g*_ are given a prior 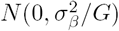 where *G =* 12,317 is the number of genes in the analysis. Integrating over *β*_*g*_ we have

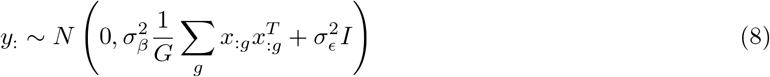

We optimize this model wrt *σ*_*β*_ and *σ*_*e*_ to obtain an estimate 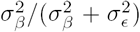 of the percent variance of *y* explained by *x*.

### Code and data availability

All the custom analysis scripts used for this project are available at https://github.com/davidaknowles/dox. The suez response eQTL mapping R package is available at https://github.com/davidaknowles/suez. The following data are available as Supplementary Data: 1) differential expression cluster assignments, 2) significant (5% FDR) eQTLs and sQTLs, 3) differential splicing results, 4) levels of cardiac troponin and the predicted transcriptomic response. In addition to the Supplementary Data included with this paper full results are hosted at http://web.stanford.edu/~dak33/dox/ including 1) gene-by-sample matrix of RNA-seq quantification (log counts per million), 2) LeafCutter intron excision quantification 3) *p-*values for all tested eQTLs, reQTLs, sQTLs, and rsQTLs. The RNA-seq FASTQ files will be added to the dbGaP database^50^ under dbGaP accession phs000185 (https://www.ncbi.nlm.nih.gov/projects/gap/cgi-bin/study.cgi?study_id=phs000185). The genotype data files cannot be shared because releasing genotype data from a subset of individuals in the pedigree would enable the reconstruction of genotypes of other members of the pedigree, which would violate the original protocol approved by the research ethics board^17^. The summary statistics for the ACT GWAS were given to us by the authors of the study^5^.

## Acknowledgements

The idea for the study was developed through conversations with Julian Solway (University of Chicago). We gratefully acknowledge Bryan Schneider for providing GWAS summary statistics. This work was supported by the Howard Hughes Medical Institute, and the US National Institutes of Health (NIH grants HL092206, HG008140, HG009431). CKB was supported by a NIH (http://www.ncats.nih.gov/ctsa) Clinical and Translational Science Award 5 TL1 TR 432-7 pre-doctoral fellowship.

## Author Contributions

YG, CKB and CO conceived the project and provided source material. CKB performed the experiments with assistance from KMP. DAK developed algorithms, and analyzed the data with assistance from JDB. JKP, CO and YG supervised the project. DAK wrote the paper with input from all authors.

## Supplementary material

**Figure S1:**
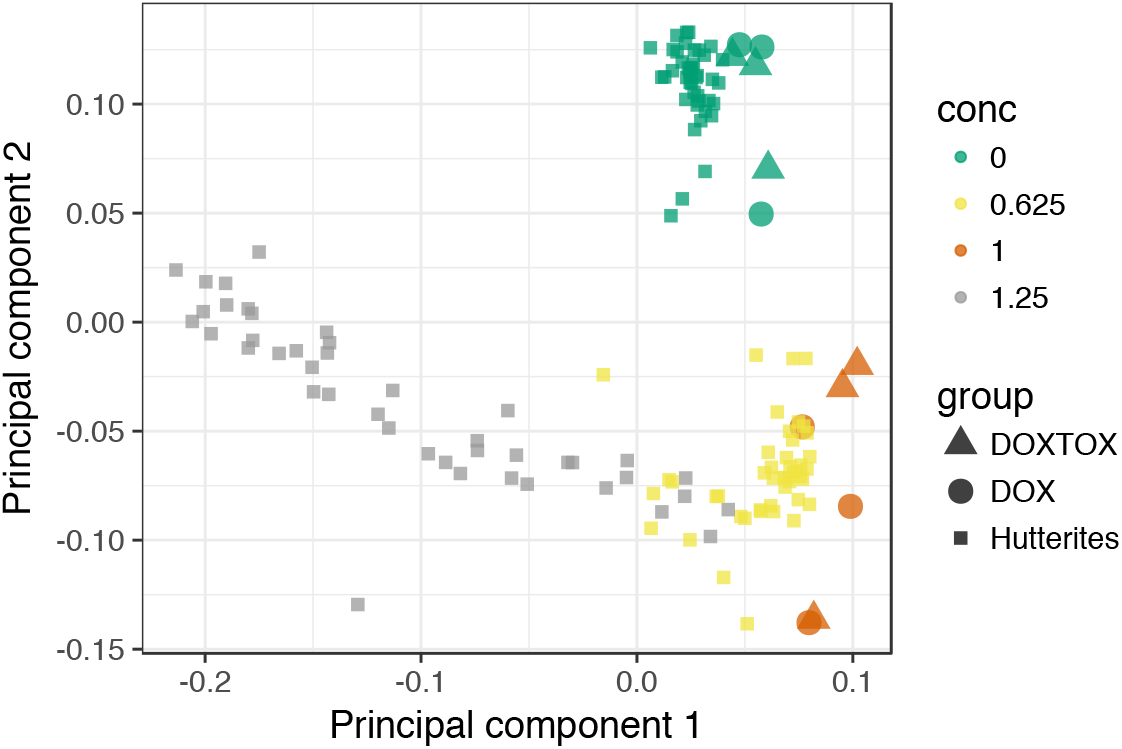
Our expression data is concordant with an existing small RNA-seq dataset^12^. DOXTOX and DOX correspond to samples from patients that did and did not develop ACT after doxorubicin chemotherapy respectively. We additionally see that the transcriptional response at higher concentrations cannot be extrapolated from that at lower concentrations. Higher concentrations are not shown since including these compresses the first PC obscuring the relevant variation.

**Figure S2:**
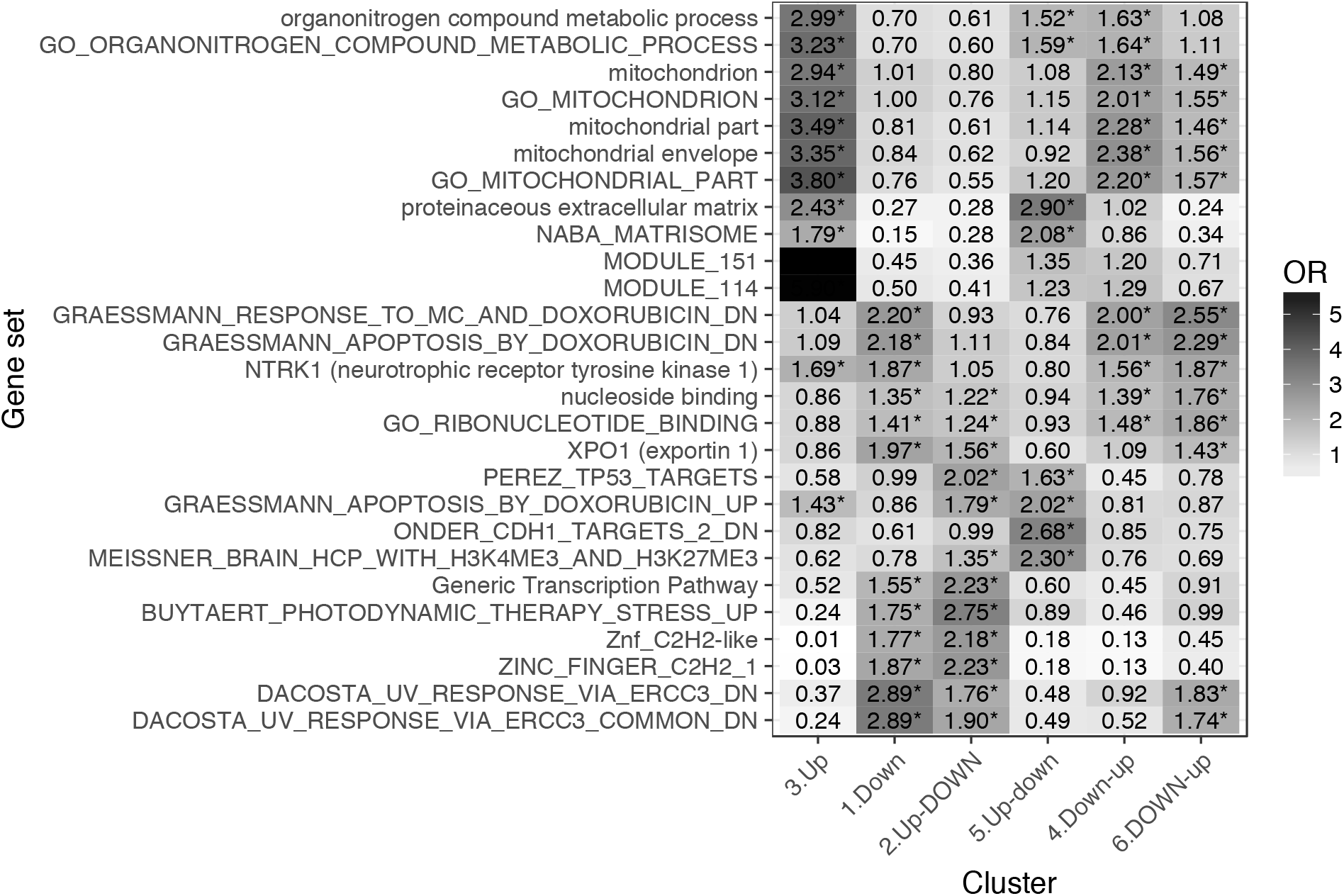
Gene set enrichment analysis of genes in each response cluster confirms expected patterns: metabolic, mitochrondrial and DNA damage processes, as well as existing doxorubicin response genes.

**Figure S3:**
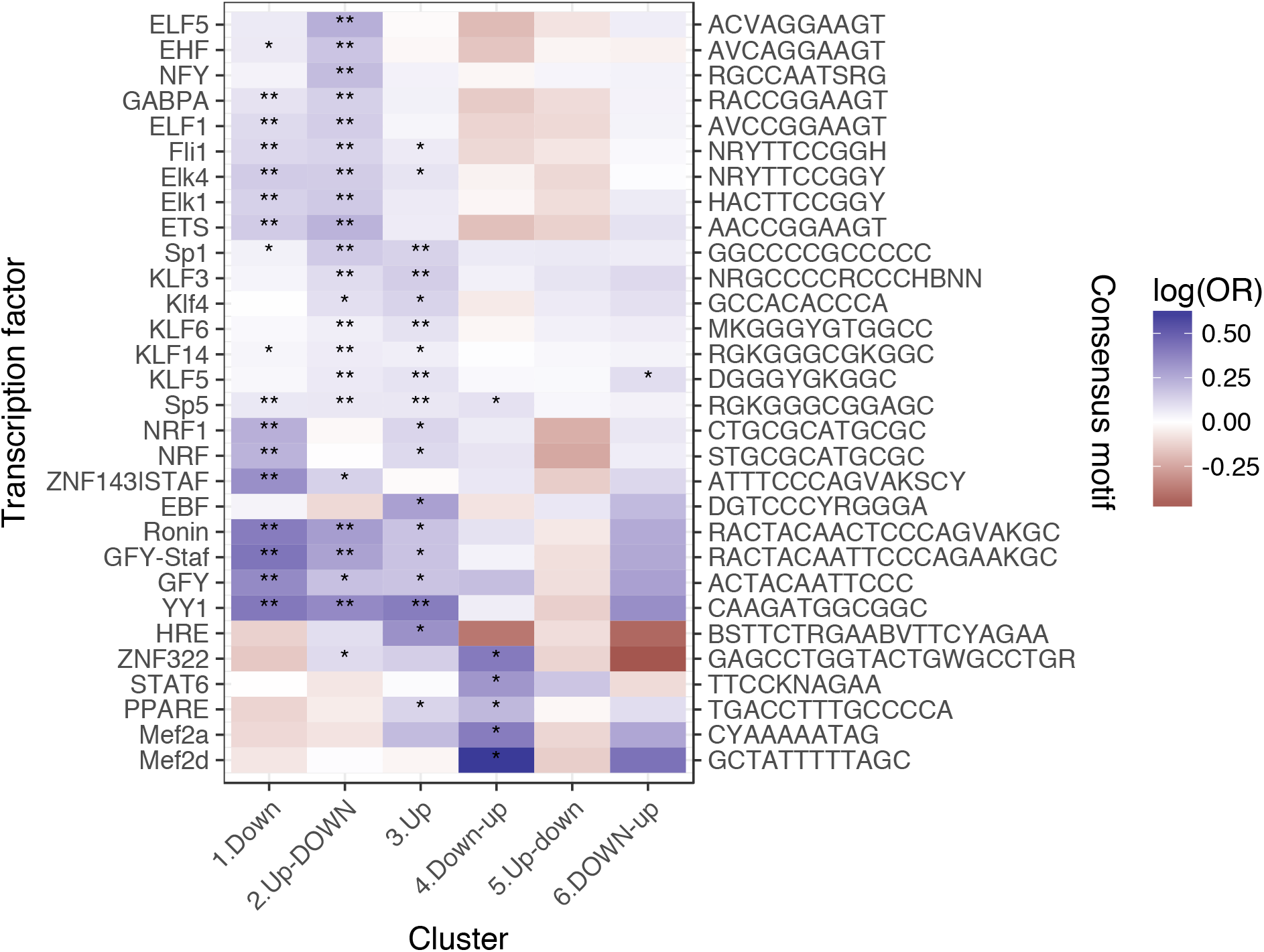
Enrichment of transcription factor binding motifs for each response pattern, using HOMER. ** denotes q < 0.05, * denotes q < 0.5.

**Figure S4:**
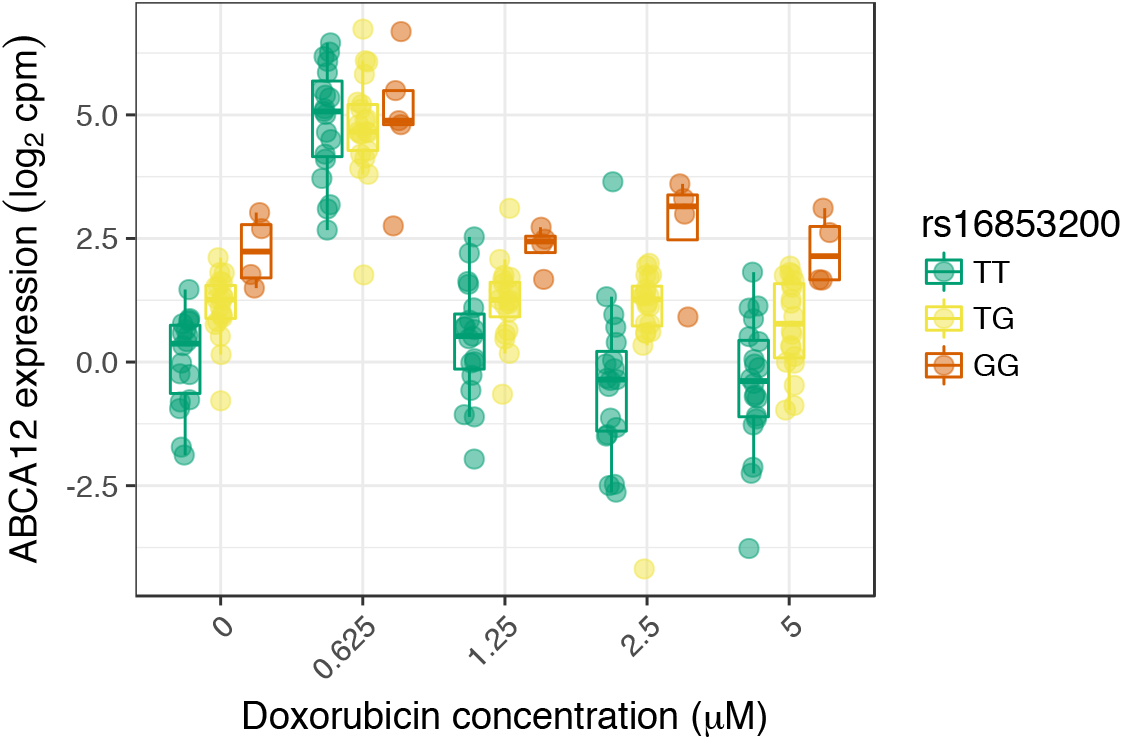
An example response expression QTL that may act through buffering at high expression levels.

**Figure S5:**
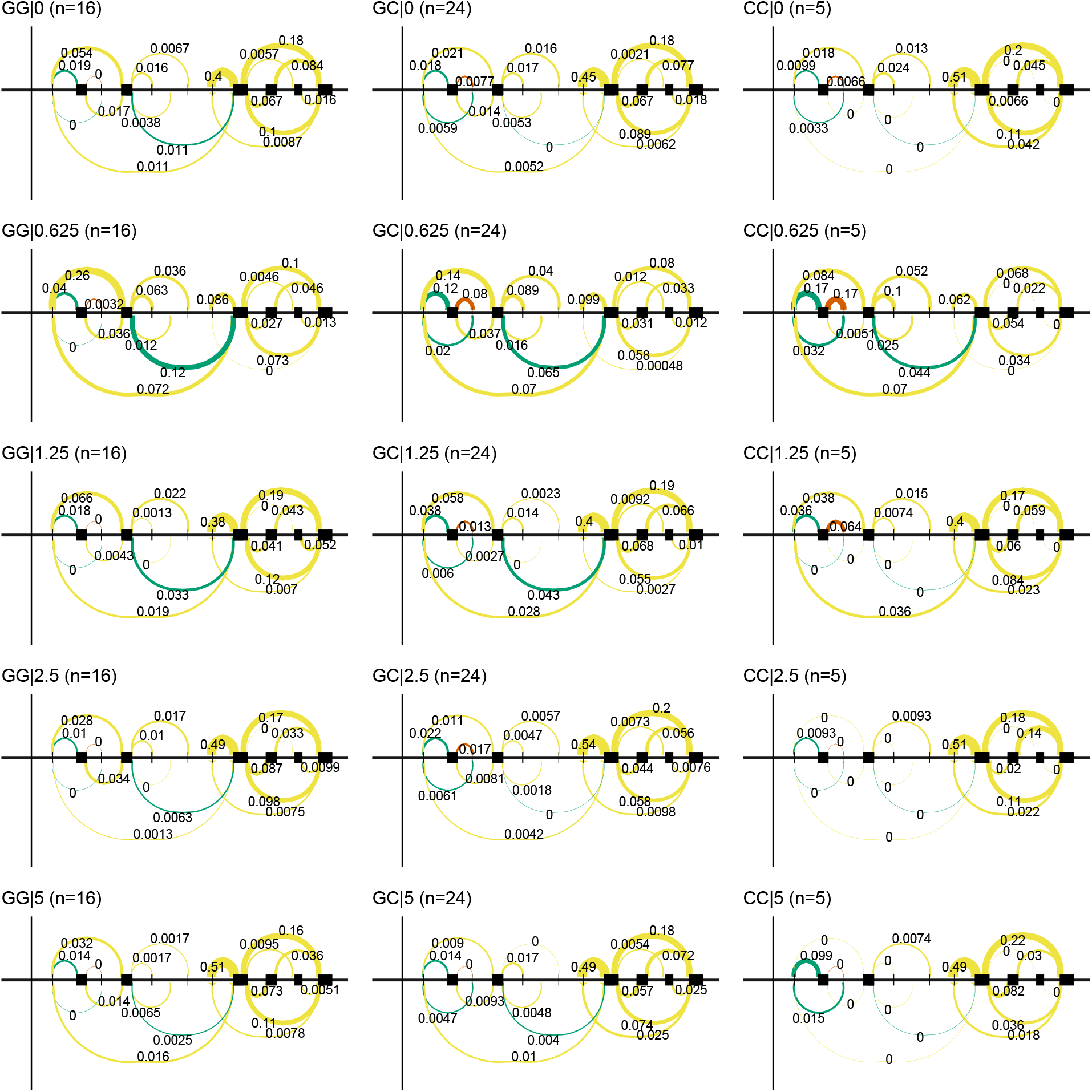
Doxorubicin exposure results in the use of a downstream alternative TSS which uncovers an association between rs896853 genotype and inclusion of the exon at chr8:94975318-94975415.

**Figure S6:**
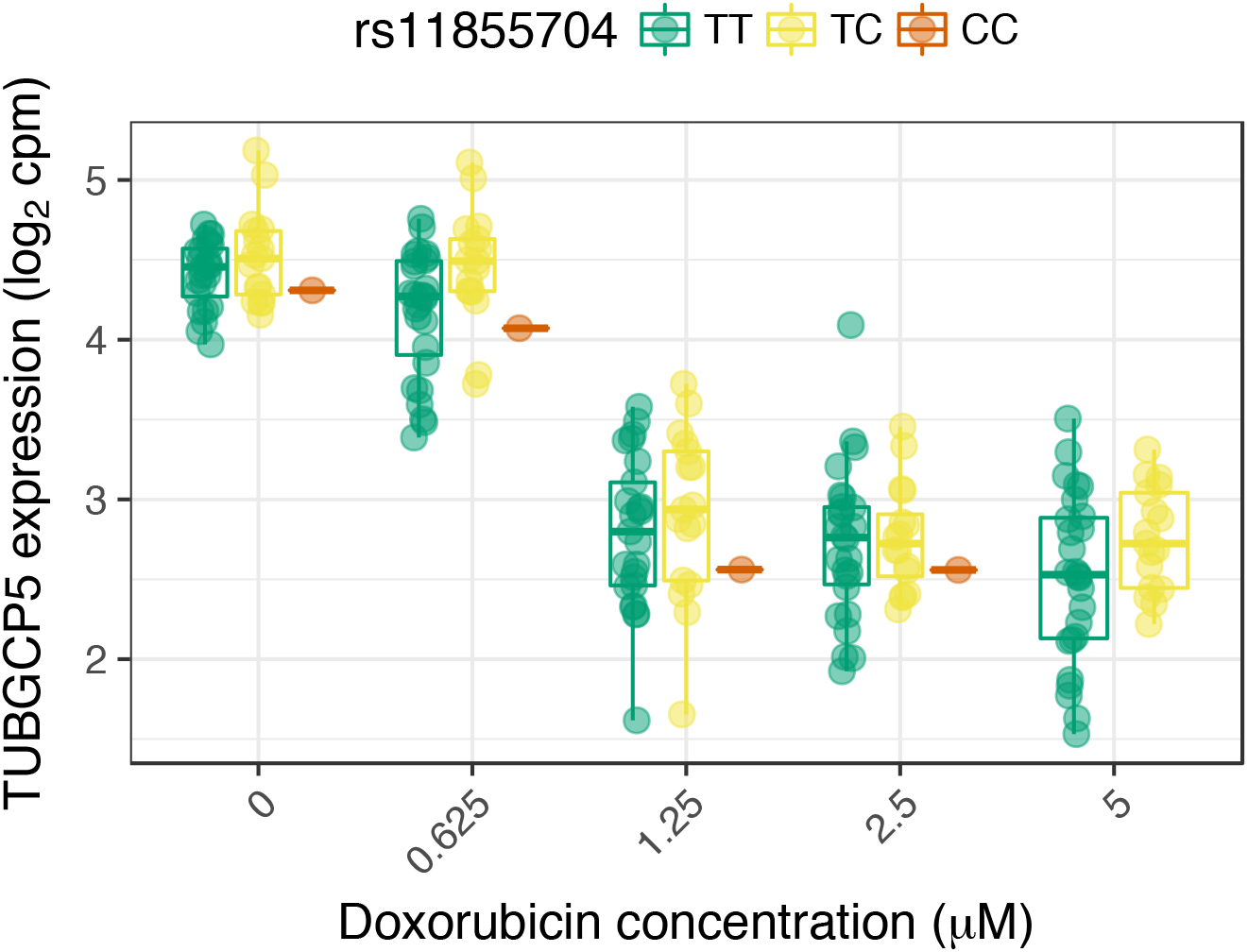
rs28714259, the one replicated variant from Schneider et al.^5^ is in high LD (R^2^ = 0.98) with rs11855704, which is a nominally significant (p = 0.036) eQTL for TUBGCP5. TUBGCP5 is strongly down-regulated in the presence of doxorubicin.

**Figure S7:**
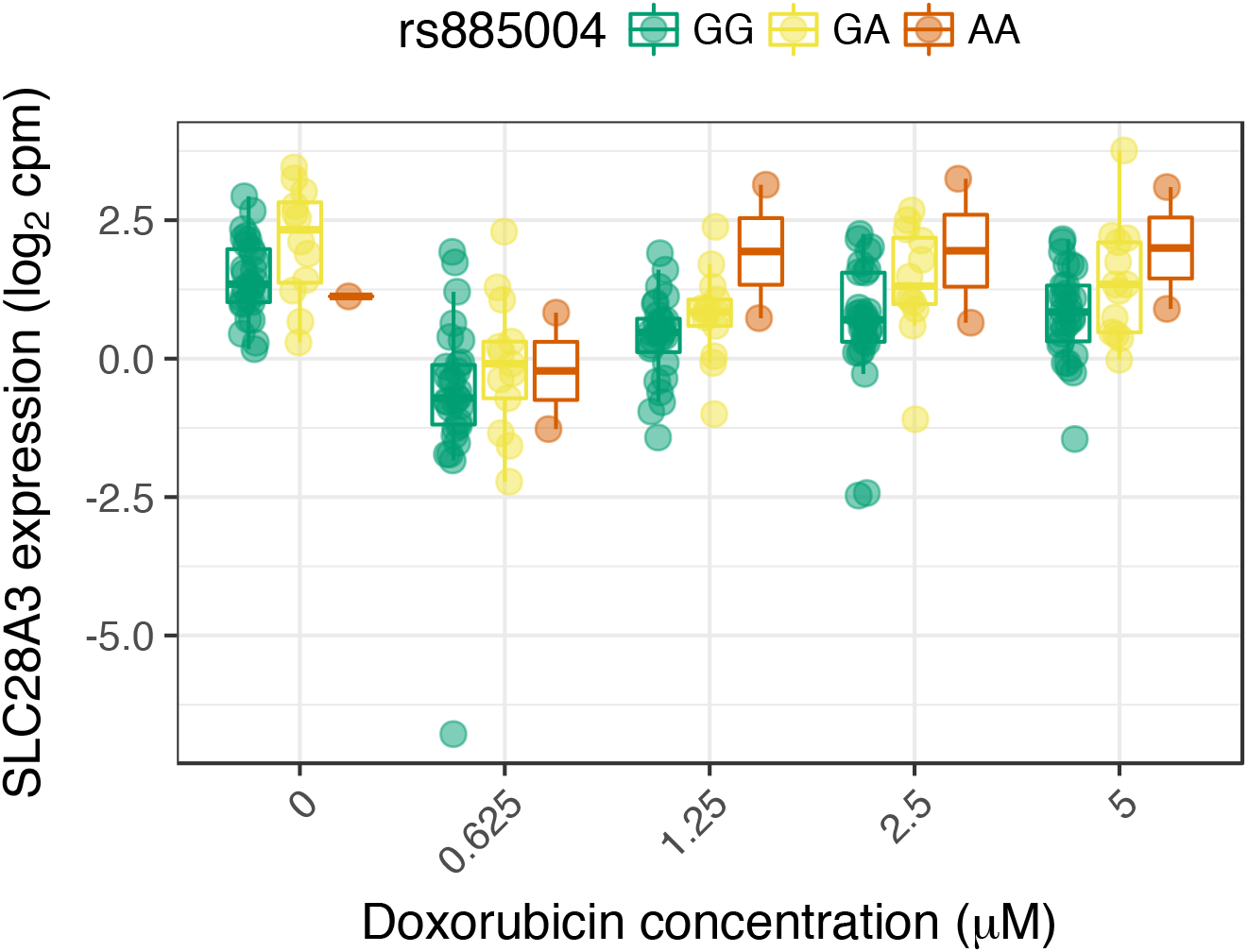
The intronic (to SLC28A3) rs885004, and the closely linked (R^2^ = 0.98) synonymous variant rs7853758, have been associated with ACT by candidate gene studies^30^,^29^. We find rs885004, but not rs7853758, has a significant marginal effect on SLC28A3 expression.

